# The impact of surveillance and control on highly pathogenic avian inuenza outbreaks in poultry in Dhaka division, Bangladesh

**DOI:** 10.1101/193177

**Authors:** Edward M. Hill, Thomas House, Madhur S. Dhingra, Wantanee Kalpravidh, Subhash Morzaria, Muzaffar G. Osmani, Eric Brum, Mat Yamage, Md. A. Kalam, Diann J. Prosser, John Y. Takekawa, Xiangming Xiao, Marius Gilbert, Michael J. Tildesley

## Abstract

In Bangladesh, the poultry industry is an economically and socially important sector, but it is persistently threatened by the effects of H5N1 highly pathogenic avian influenza. Thus, identifying the optimal control policy in response to an emerging disease outbreak is a key challenge for policy-makers. To inform this aim, a common approach is to carry out simulation studies comparing plausible strategies, while accounting for known capacity restrictions. In this study we perform simulations of a previously developed H5N1 influenza transmission model framework, fitted to two separate historical outbreaks, to assess specific control objectives related to the burden or duration of H5N1 outbreaks among poultry farms in the Dhaka division of Bangladesh. In particular, we explore the optimal implementation of ring culling, ring vaccination and active surveillance measures when presuming disease transmission predominately occurs from premises-to-premises, versus a setting requiring the inclusion of external factors. Additionally, we determine the sensitivity of the management actions under consideration to differing levels of capacity constraints and outbreaks with disparate transmission dynamics. While we find that reactive culling and vaccination policies should pay close attention to these factors to ensure intervention targeting is optimised, across multiple settings the top performing control action amongst those under consideration were targeted proactive surveillance schemes. Our findings may advise the type of control measure, plus its intensity, that could potentially be applied in the event of a developing outbreak of H5N1 amongst originally H5N1 virus-free commercially-reared poultry in the Dhaka division of Bangladesh.

## Introduction

Influenza is a respiratory infection of mammals and birds caused by an RNA virus in the family of Orthomyxoviridae [1]. There are four types of influenza viruses: A, B, C and D. Of these four types, the zoonotic capability of influenza A viruses makes them the most significant in an epidemiological and public health context, being associated with most of the widespread seasonal influenza epidemics and the type of influenza capable of causing occasional global pandemics. While the natural hosts of influenza A viruses are aquatic bird species, these viruses occasionally spillover into other animal hosts, including domestic poultry, pigs, horses, a variety of carnivores and marine mammals [2]. Sporadically, the viruses adapt to their new animal hosts, leading to enzootic virus circulation for sustained periods. However, apart from a few cases of reputed direct zoonotic transmission of influenza A viruses to humans from wild birds, due to close contact and de-feathering activities [3, 4], humans have been primarily infected with zoonotic influenza viruses via intermediate species to which human exposure is more frequent. Domestic livestock such as pigs and poultry have a key role in this regard. Accordingly, influenza A is not considered an eradicable disease, rather prevention and control are the only realistic goals [5].

The prevention and control of Highly Pathogenic Avian Influenza (HPAI) in poultry is of paramount importance, with HPAI viruses causing severe disease in domestic poultry with a high death rate [6]. The specific intervention actions to be taken with regards to regulating live bird markets (LBMs), imposing movement restrictions or quarantine measures, and culling and vaccinating can vary according to local circumstances and from country to country. A single solution for all situations is unattainable, and a balance must be established among effective, feasible and socially acceptable control measures that safeguard the short-term and long-term livelihoods of farmers and the health of the population.

In general, however, a number of basic measures are common to all circumstances. One such measure is that infected birds and those in contact with them must be humanely and safely culled to halt spread of the disease. Humane culling limits spread of disease by decreasing the amount of virus shed from any one site. However, culling alone usually cannot completely prevent further spread because some virus will have been released before culling commences, and often before the disease is detected. As a result, pre-emptive culling (the culling of animals before they are found to be infected) can be used to attempt to make this a more proactive measure. Use of widespread pre-emptive culling based on defined areas around an outbreak has been a standard implementation of this protocol [7].

Disease control programs may also aim to create impediments to spread. An essential part of creating impediments is to establish an environment in which there are relatively few locations that could become easily infected, with vaccination one of the main methods available for achieving such a goal [7]. Vaccination against HPAI aims to prevent clinical disease as well as to reduce levels of virus shed into the environment and stop infection spreading. In parts of Asia, vaccination programs have been implemented and encouraged as part of a control program in poultry. Vietnam is a notable case, with it being found that within-flock reproductive numbers - i.e. the expected number of secondary cases from an average primary case in an entirely susceptible population - for premises reporting H5N1 infection were lower in an outbreak period using both depopulation and nationwide systematic vaccination campaigns, compared to an outbreak period employing depopulation control measures alone [8]. Recent positive developments have seen vaccines against H5N1 and H7N9 prevent oral and faecal viral shedding, thus stopping transmission from one bird to another [9]. Of particular importance is ensuring the vaccines used have high efficacy. In Bangladesh, vaccines against HPAI have been available since 2012 for use on commercial layer and breeder farms (M.G. Osmani and M.A. Kalam, personal communication). However, a recent H5N1 surveillance study found that anti-H5 seropositivity levels were similarly low in vaccinated and unvaccinated chickens, suggesting that the vaccine is not effective in inducing protective antibody levels against H5N1 and, as a result, demonstrating a need for updated poultry vaccines [10].

Policy effectiveness will depend critically on how swiftly clinical cases are diagnosed and the speed with which the chosen control measure can be administered. By employing active surveillance of premises (i.e. activities that are frequent, intensive and aim at establishing the presence or absence of a specific disease), the time for identifying cases and notifying officials of an infected flock may be reduced.

Although active surveillance activities can be expensive and time-consuming, there are notable examples of the benefits of strengthening influenza surveillance programs. Intensification of surveillance has helped control and limit the spread of HPAI viruses among poultry on a national scale (e.g. Nigeria [11]), while early detection of HPAI H5N1 viruses through enhanced surveillance in wild birds and domestic poultry has been a key measure to ensure rapid disease control on a continental scale in the case of the European Union [12]. Improved influenza virus surveillance in pigs revealed that influenza virus transmission from humans to swine is far more frequent than swine-to-human zoonosis [13]. The public availability of genetic sequence data from databases such as GenBank has allowed pioneering studies to come into fruition, setting out to characterise the cross-species nature and the migration of influenza A viruses on a global scale [14]. In addition, there are probable long-term advantages to be gained from active surveillance which outweigh the costs. These advantages include trade benefits, with eventual proof of disease absence allowing the opening-up of hitherto untapped markets. Further, for diseases such as rinderpest, beginning active surveillance meant vaccination could cease, saving sizeable amounts of money that otherwise would have been spent on blanket vaccination campaigns [15].

In conjunction with this collection of possible actions, distinct stakeholders may have disparate control objectives. As a consequence, stakeholders may have different metrics of management success that they are most interested in optimising. Crucially, alternative objectives may require differing approaches to ensure outcomes are optimal [16, 17]. Objectives may only depend upon a single, measurable outbreak burden quantity, such as duration. On the other hand, objectives may be linked to multiple outbreak quantities and be treated in monetary terms, as has been previously seen in the context of other livestock diseases such as foot-and-mouth disease [17, 18]. Throughout this paper, we concentrate on the former category of objectives, namely through the following outbreak burden facets: duration, size (in the form of total number of premises infected during the course of the outbreak), cumulative number of poultry culled and spatial extent. Whilst a rigorous cost analysis is beyond the scope of this paper, the application (to this setting) of objective functions that are treated in monetary terms is an avenue for future work.

We focus in this study on commercial poultry premises in Bangladesh. In addition to being a country that has suffered from recurrent H5N1 outbreaks in commercial poultry as recently as 2012 [19], with H5N1 viruses now considered to be endemic in the nation [20, 21], Bangladesh is a prime candidate for being the source of newly emerging influenza strains with pandemic-causing potential. The reasons for this are twofold: first, Bangladesh is one of the most densely populated countries in the world [22]; second, Bangladesh already has a substantial poultry population (1194 birds/km^2^) and the poultry industry is going through a period of rapid intensification [23]. The aforementioned factors are underlined by the recent emergence of a new genotype of HPAI H5N1 viruses in the country that are now dominant and represent the current threat to domestic poultry and humans in the region [24].

Yet, recently conducted endemicity studies in two major poultry producing divisions of Bangladesh did not yield H5 positives from any of the commercial farms sampled (E. Brum, unpublished observations). These findings indicate that the commercial farms in major poultry producing areas are managing to stay free from H5 HPAI, while an operational goal of a prospective strategy for control of H5N1 HPAI in Bangladesh is to protect poultry in farms and villages to decrease the prevalence of H5N1 (E. Brum and M.S. Dhingra, personal communication; see Supporting Information for further details). Under these circumstances, H5N1 infection returning to these regions may spark a larger outbreak with characteristics akin to an epidemic. For that reason, it is vital to assess the capability of various intervention approaches in curbing the burden and/or duration of H5N1 HPAI outbreaks in HPAI-free localities.

The key platforms of current HPAI control programs in Bangladesh, that are directed towards the commercial poultry industry, are focused on case detection, identification of premises deemed to be in direct contact with a premises reporting infection, and subsequent stamping out of flocks with reported infection [21]. Bangladesh has however adopted, or has the potential to implement, each of the intervention types described above. Historically, Bangladesh adopted a ring culling approach to combat HPAI outbreaks. Prior to 2008, poultry flocks within 1km of premises with confirmed HPAI infection were designated to be culled (M.A. Kalam, personal communication). Furthermore, with vaccines against HPAI now being available (since 2012) for use on commercial layer and breeder farms, ring vaccination has become an implementable control management action. In terms of active surveillance, from 2008 to 2012 a small-scale active surveillance system was run. This comprised of teams of community health workers across the country, each monitoring specified farms and reporting to livestock officers mortality events and the presence of any clinical signs of disease (M.G. Osmani and M.A. Kalam, personal communication). Thus, for Bangladesh ring culling, ring vaccination and active surveillance are representative of HPAI control policies that have been implemented historically, are currently in use or that could be pursued as management alternatives in the future (for additional details on pre-existing and prospective response protocols for the control of H5N1 HPAI amongst poultry in Bangladesh, see Supporting Information).

In this paper, we evaluate the above assortment of intervention styles in opposing outbreaks of H5N1 HPAI among commercial poultry premsises within the Dhaka division, Bangladesh. These assessments are performed in the context of commercial poultry premises in region beginning free of H5N1 HPAI. We also explore the potential impact these measures could have if capacities for enacting control increase over the current capacity. Assessments were conducted with respect to optimising particular control objectives that were dependent upon measurable outbreak burden quantities (such as outbreak size and duration). This analysis was done via simulations of our H5N1 influenza transmission model that has previously been fitted to outbreak data in the Dhaka division [25], allowing the optimisation of decision making under uncertainty in a principled way. Specifically, we aimed to ascertain both the required intensity of culling and vaccination measures, and type of active surveillance scheme, to optimise a given control objective. Our three primary focuses were then as follows: (i) analyse variability in these choices if in a setting where transmission is believed to be predominately from premises-to-premises, versus the scenario where importations and other external environmental/ecological factors are also considered; (ii) inform decisions regarding intervention prioritisation and implementation when under resource constraints that limit control capacity; (iii) determine the sensitivity of the choice of management action to epidemiological characteristics, by considering outbreaks with disparate transmission dynamics.

## Methods

### The data

Throughout 2010, the Bangladesh office of the Food and Agriculture Organisation of the United Nations (FAO/UN) undertook a census of all commercial poultry premises, listing 65,451 premises in total, of which 2,187 were LBMs. Each premises was visited once, recording location and the number of avian livestock present during the visit within these categories: layer chickens, broiler chickens, ducks, others (e.g. turkeys, quails). Within the census data there were instances of multiple premises having the same location (i.e. identical latitude and longitude co-ordinates). For these occurrences the avian livestock populations were amalgamated, giving a single population for each category at each location.

Of the non-market locations, 23,412 premises had blank entries for all avian types. Blank entries corresponded to no poultry being present when the census visit occurred, due to the premises either being between poultry stocks or being temporarily closed by the farmer due to an ownership transfer taking place, rather than data entry errors (M.G. Osmani, personal communication). We made a simplifying assumption that at any given time an equivalent proportion of premises would not have any avian livestock at the premises. Therefore, we did not make use of these locations in our analysis. While not discussed here, the sensitivity of model outputs to this assumption requires further consideration.

Owing to the small number of premises in the Dhaka division recorded as having ducks or other poultry types present (around 20), our model simulations comprised purely those premises recorded as having layer and/or broiler chicken flocks present. This totalled 13,330 premises.

Between 2007 and 2012, there were six epidemic waves of H5N1 among poultry in Bangladesh, resulting in a total of 554 premises with confirmed infection and over 2,500,000 birds being destroyed. In previous work [25], we developed a suite of nested models for the Dhaka division that were fitted to the two largest epidemic waves, wave 2 (September 2007 to May 2008) and wave 5 (January 2011 to May 2011), resulting in a total of 232 and 161 premises becoming infected, respectively (see Supporting Information for further epidemiological data details). In cases where there were discrepancies between flock size from the poultry case dataset and the 2010 census, we defaulted to the poultry case dataset.

### Mathematical model for H5N1 transmission

In this paper, we utilise a previously developed model framework [25] and investigate the impact of a range of control and surveillance strategies on different control objectives when there is uncertainty about epidemic dynamics and resource capacity. The model is a discrete-time compartmental model, where the individual poultry premises is the epidemiological unit of interest. Consequently, layer and broiler flock sizes at each premises were combined to give an overall poultry population. At any given point in time a premises *i* could be in one of four states, *S, I, Rep* or *C*: *i* ∈ *S* implies premises *i* was susceptible to the disease; *i* ∈ *I* implies premises *i* was infectious and not yet reported; *i* ∈ *Rep* implies premises *i* was still infectious, but had been reported; *i* ∈ *C* implies that premises *i* had been culled. In other words, all poultry types within a premises become rapidly infected such that the entire premises can be classified as Susceptible (*S*1), Infected (*I*), Reported (*Rep*) or Culled (*C*).

The reporting delay, time taken for a premises to transition from state *I* to *Rep*, accounts for a premises being infectious before clinical signs of H5N1 infection are observed, which may not be immediate [26], followed by the time taken for premises owners to notify the relevant authorities [21]. While the poultry epidemic was ongoing we assumed a premises was not repopulated once culled.

The force of infection against a susceptible premises *i* on day *t* was comprised of two terms: (i) the force of infection generated by an infectious premises *j* (*η_ij_*), (ii) a ‘spark’ term (*ϵ*_*j*_) to allow for spontaneous, non-distance dependent infections that were unexplained by the susceptibility, transmissibility and kernel components of the model [27]. This captures factors such as importations from outside the study region and transmission from virus-contaminated environments (i.e. fomites). Further, despite backyard poultry not being explicitly included within these models its contribution to the force of infection could be incorporated into *ϵ*_*j*_.

As a result, the total force of infection has the following general form:

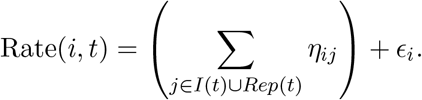

We assume a seven day delay from infection to reporting (unless specified otherwise), in line with the results of previous work [25, 28]. The contribution by infected premises *j* to the force of infection against a susceptible premises *i* satisfies

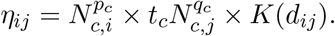

*N*_*c,i*_ is the total number of chickens recorded as being on premises *i*, *t_c_* measures the individual chicken transmissibility, *d_ij_* is the distance between premises *i* and *j* in kilometres, and *K* is the transmission kernel to capture how the relative likelihood of infection varies with distance. The model also incorporated power law exponents on the susceptible population, *p_c_*, and infected population, *q_c_*. Including power law exponents allows for a non-linear increase in susceptibility and transmissibility with farm size, which has previously been shown to provide a more accurate prediction of farm-level epidemic dynamics [29].

The transmission kernel *K* in our model is Pareto distributed such that:

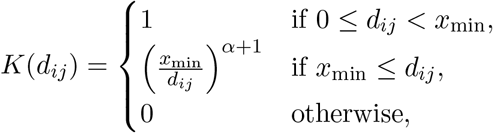

where *x*_min_ is the minimum possible value of the function (set to 0.1, corresponding to 100 metres, with all between location distances less than 100 metres taking the 100 metre kernel value) and *α* ≥ −1. Values of *α* close to −1 give a relatively constant kernel over all distances, with *α* = −1 corresponding to transmission risk being independent of distance. As *α* increases away from −1 localised transmission is favoured, with long-range transmission diminished.

The spark term was the same fixed value for every premises, *ϵ*, with the total rate of infection against a susceptible premises *i* on day *t* satisfying

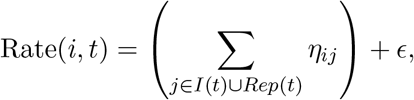

The previous model fitting study found the wave 5 division-level model, compared to the wave 2 fitted model, had a stronger preference for short-range transmission, with the flock size of infectious premises also having a more prominent role in the force of infection [25]. This allowed us to explore the sensitivity of the management actions under consideration to epidemics with disparate transmission dynamics. Complete listings of the inferred parameter distributions for both models are provided in Table S2.

### Poultry control policies of interest

In the event of outbreaks of H5N1, a range of policies may be implemented to reduce the risk of further spread of disease. We investigated the relative effect of the implementation of three poultry-targeted policy actions, ring culling, ring vaccination and active surveillance, which are representative of controls that have been implemented historically, are currently in use or that could be pursued as management alternatives in the future in Bangladesh.

There are often restrictions on the resources available for enforcing such interventions, limiting the number of poultry and/or premises that can be targeted on any given day. As a consequence, we imposed daily capacities on the maximum number of poultry and the maximum number of premises targeted by each control action, with three differing levels of severity related to the availability of resources.

We investigate here resource constraints that are representative of current capacities to enact control measures in Bangladesh, but in addition explore the potential impact of interventions should capacities be larger than are currently the case in the country. By examining a range of constraints, we could establish if the action determined optimal was sensitive to the daily capacity to carry out control. Resource limits exceeding the upper capacity levels considered here were not investigated due to requiring a longer-term build up of government resources to be attainable (M.G. Osmani, personal communication).

In each case a baseline control measure of only culling reported premises was performed, with premises being culled on the same day they were reported if possible (with respect to the resource constraints in place). Note that culling of premises reporting infection was carried out in all subsequent control strategies outlined below.

### Ring culling

For this choice of action, in addition to the culling of premises reporting infection, all premises within a specified distance of locations with confirmed infection were marked for culling. The distances evaluated here ranged from 1-10km (in 1km increments). In order to simulate the effect of differing resource constraints within the Dhaka division, we imposed three conditions, based upon low, medium and high culling capacities (see Table 1 for a listing of tested capacity values).

**Table 1:** Limits for carrying out the specified control option at low, medium and high capacity levels.

| Control strategy | Capacity level | Bird limit (per day) | Premises limits |  |  |
| --- | --- | --- | --- | --- | --- |
|  |  |  | Per day | Per outbreak | Coverage |
| Ring cull | Low | 20,000 | 20 |  |  |
|  | Medium | 50,000 | 50 |  |  |
|  | High | 100,000 | 100 |  |  |
| Ring vaccination | Low | 20,000 | 20 |  |  |
|  | Medium | 50,000 | 50 |  |  |
|  | High | 100,000 | 100 |  |  |
| Reactive surveillance | Low |  |  | 25 |  |
|  | Medium |  |  | 50 |  |
|  | High |  |  | 100 |  |
| Proactive surveillance | Low |  |  |  | 5% |
|  | Medium |  |  |  | 10% |
|  | High |  |  |  | 25% |

Shaded cells indicate limit classes that were not applicable under the given type of control strategy.

To clarify, premises reporting infection were prioritised above all others for culling, ordered by the date of reporting. For those premises designated for ring culling that were not infected, the order of priority was determined using a distance-based approach, with resources allocated from the outer edge and moving inwards to the centre (an ‘outside-to-centre’ approach). In other words, following the determination of premises situated within the ring established around a premises reporting infection, distances between all such premises and the infected premises were computed with the premises then culled in descending distance order. Note that all premises in the ring established around the initially reported infected premises had to be treated before moving on to locations that were contained within rings established around the next set of subsequently reported infected premises.

### Ring vaccination

For this choice of action, all premises within a specified distance of any premises reporting infection were listed for vaccination. As with ring culling, the ring radii evaluated ranged from 1-10km (in 1km increments). In light of previous research highlighting apparent discrepancies between the vaccine strain and the viruses in circulation in Bangladesh [10] we did not assume perfect vaccine efficacy, but instead set efficacy to 70% (unless specified otherwise). While this efficacy is not guaranteed to fully agree with the true efficacy of currently administered vaccines, it considers a general situation where the proposed vaccine possesses a reasonable capability to suppress the circulating strain. We assumed a baseline effectiveness delay value of seven days to account for the time required for suitable immune protection to develop after the vaccine was administered (M.G. Osmani and M.A. Kalam, personal communication). With the epidemiological unit of interest being the individual poultry premises, we assumed for successfully vaccinated flocks (i.e. vaccinated premises that did not become infected during the post-vaccination effectiveness delay period) that, given a 70% vaccine efficacy, 30% of the flock remained susceptible to infection (and as a consequence able to transmit infection).

As the vaccination strategies considered here also involved the culling of reported premises, we had to make an assumption regarding how culling and vaccination aspects should be factored into the resource limits. We were informed that while culling would be carried out by DLS (Department of Livestock Services) staff, vaccines would be administered by the farms themselves under the supervision of DLS staff (M.A. Kalam, personal communication). Accordingly, we treated culling and vaccination activities as being independent of each other, assigning separate resource limitations to each control action. As for ring culling, low, medium and high capacity settings were investigated (Table 1).

There was no limit on the cumulative number of vaccine doses available. An outside-to-centre resource allocation prioritisation approach was used for vaccination, matching the ring culling prioritisation procedure.

### Active Surveillance

The active surveillance actions of interest here concentrated on the earlier detection of clinical signs of disease within poultry flocks. In model simulations of active surveillance initiatives, premises undergoing active surveillance had their notification delay reduced from seven to two days. A two day delay was chosen, and not a larger reduction to a single day or the complete removal of the reporting delay, to align with the shortest delay in detecting clinical signs that is realistically attainable under ideal conditions. Such a presumption has been made in prior studies [30], and accounts for the fact that a flock can be infectious before clinical signs of H5N1 infection are observed, which may not be immediate even when active surveillance procedures are in place [26]. Note that there were no other control actions in place beyond this and the culling of flocks at premises reporting infection (which abided by the previously discussed capacity limitations).

Four active surveillance strategies were compared based on two distinct types of implementation. The first two surveillance strategies we consider are reactive in nature. In reactive schemes, holdings undergo active surveillance if within a given distance of premises reporting infection. We imposed a limit on the number of premises that could be treated in this way. Thus, when resource thresholds were exceeded, only those premises deemed to be of higher priority underwent active surveillance, with the following two prioritisation strategies studied: (i) ‘reactive by distance’, with premises ordered by distance to the focal premises, nearest first (i.e. inside-to-out approach); (ii) ‘reactive by population’, with premises ordered in descending flock size order. For both schemes the ring size for active surveillance was set at 500m.

The next two surveillance strategies under consideration are proactive approaches, with a specified proportion of premises within the Dhaka division selected by some designated criteria to undergo constant active surveillance. The two criteria evaluated here were: (i) ‘proactive by population’, by ranking all premises in descending flock size order, (ii) ‘proactive by premises density’, by ranking all premises in descending order of premises density within 500m.

For both kinds of active surveillance (reactive and proactive approaches), we again considered three capacity settings (low, medium, high), with the specific limits stated in Table 1.

### Simulation outline

The simulation procedure employed here used the Sellke construction [31]. A desirable characteristic of this framework is that the inherent randomness of an epidemic realisation can be encoded at the beginning of the simulation with a random vector *Z* of Exp(1) distributed resistances. Once calculated, the resultant epidemic can be constructed from the deterministic solution of the infection process and removal (i.e. culling) times. For that reason, this method provides improved comparisons of interventions, with direct comparison of a collection of control measures achieved by matching values of *Z* at the epidemic outset. All calculations and simulations were performed with Matlab^®^.

### Choice of control policy based on outbreak origin

For this series of simulations we were interested in elucidating the intensity of control actions necessary to minimise epidemic severity based on the district of outbreak origin, and how this differed between the two fitted models with their contrasting premises-to-premises transmission dynamics. To be able to ascertain the true impact of outbreak origin on the epidemic outcomes of interest we assumed premises infection was predominately driven by premises-to-premises transmission, with no infection of premises arising due to external factors. As a consequence, the background spark term *ϵ* was set to zero in all runs, while in each run an initial cluster of three infected premises was seeded in one of the 18 districts situated within the division (see Fig. 1).

**Fig. 1.**
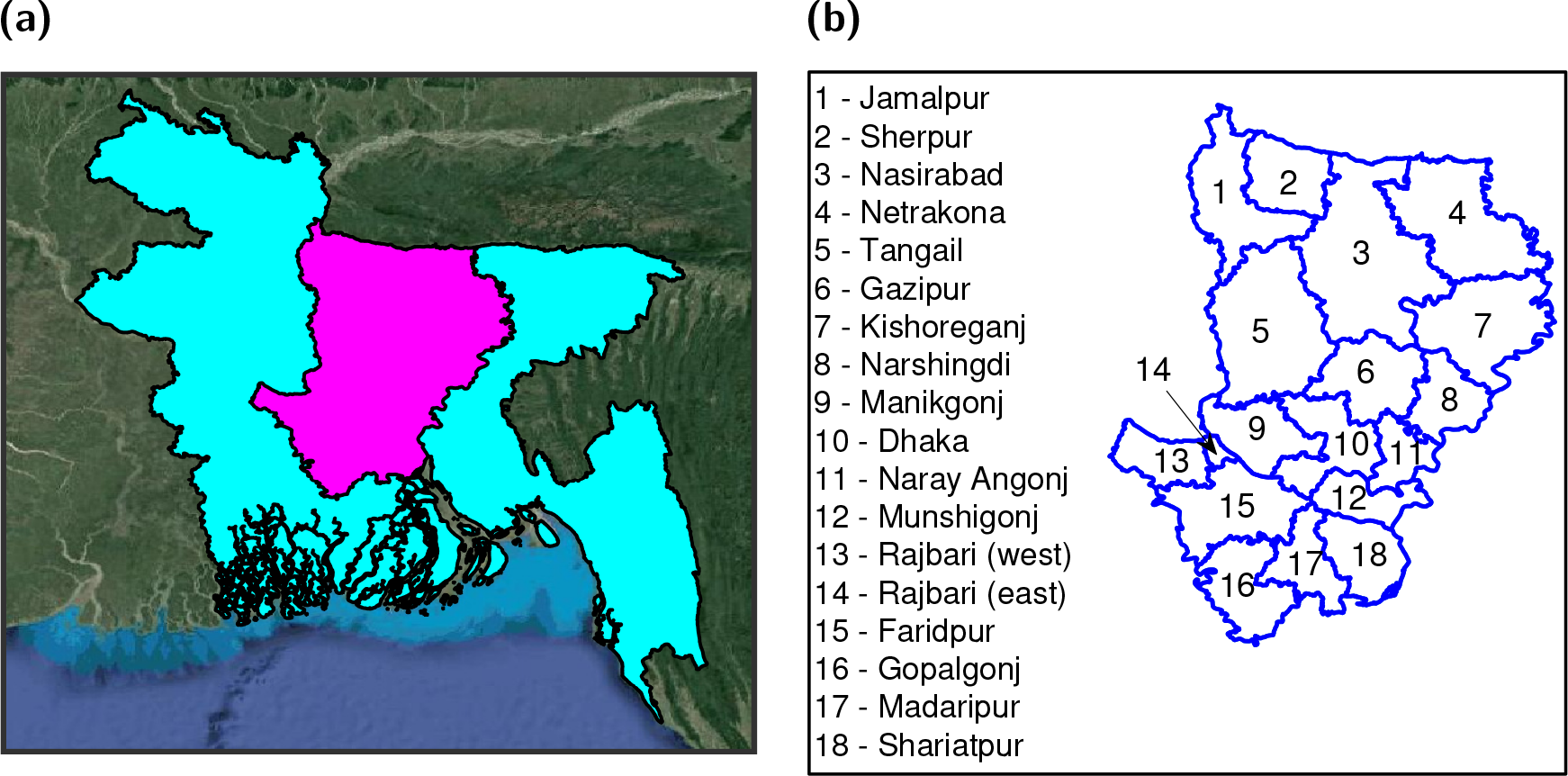
Dhaka administration region locator maps. **(a)** Locator map depicting the location of Dhaka division, shaded in magenta, within Bangladesh, shaded in cyan. **(b)** Locator map naming each district that is contained within the Dhaka division.

For each culling, vaccination, and active surveillance management action, we performed 1,000 simulation runs with the wave 2 fitted transmission model and between 500 to 1,000 simulation runs with the wave 5 fitted transmission model. A consistent set of distinct sampled parameter values (obtained previously via MCMC) and initial seed infection locations were used across these runs to aid intervention comparisons. The particular control objectives of interest here were focused on either reducing the expected length of an outbreak, or minimising the likelihood of an outbreak becoming widespread. To this end, the summary outputs analysed for this scenario were as follows: (i) mean outbreak duration, (ii) probability of an epidemic (where we subjectively define an outbreak as an epidemic if there are infected premises in five or more districts, with the total number of infected premises exceeding 15).

### Choice of control policy in presence of external factors

Our second scenario of interest was to determine the optimal control strategy when an outbreak is ongoing and infection may arise anywhere within the division, in addition to premises-to-premises transmission dynamics. Simulations for this scenario incorporated the background spark term *ϵ*, with a single initial infected premises placed anywhere in the division.

We stipulated a simulated outbreak to be complete once a specified number of consecutive infection-free days had occurred. For the wave 2 fitted model, a value of 28 days gave a simulated median epidemic length (using infected premises culling only, with reporting to culling times weighted by the empirical probability mass function) that corresponded well with the data (Fig. 2(a)). On the other hand, a 14 day period with no premises becoming infected was more suitable for the wave 5 fitted model (Fig. 2(b)), with runs using the 28 day infection-free condition giving, in general, longer outbreak periods than the observed data (Fig. 2(c)). As a consequence, the infection-free condition values were set to 28 days and 14 days for runs with the wave 2 and 5 fitted models respectively.

**Fig. 2.**
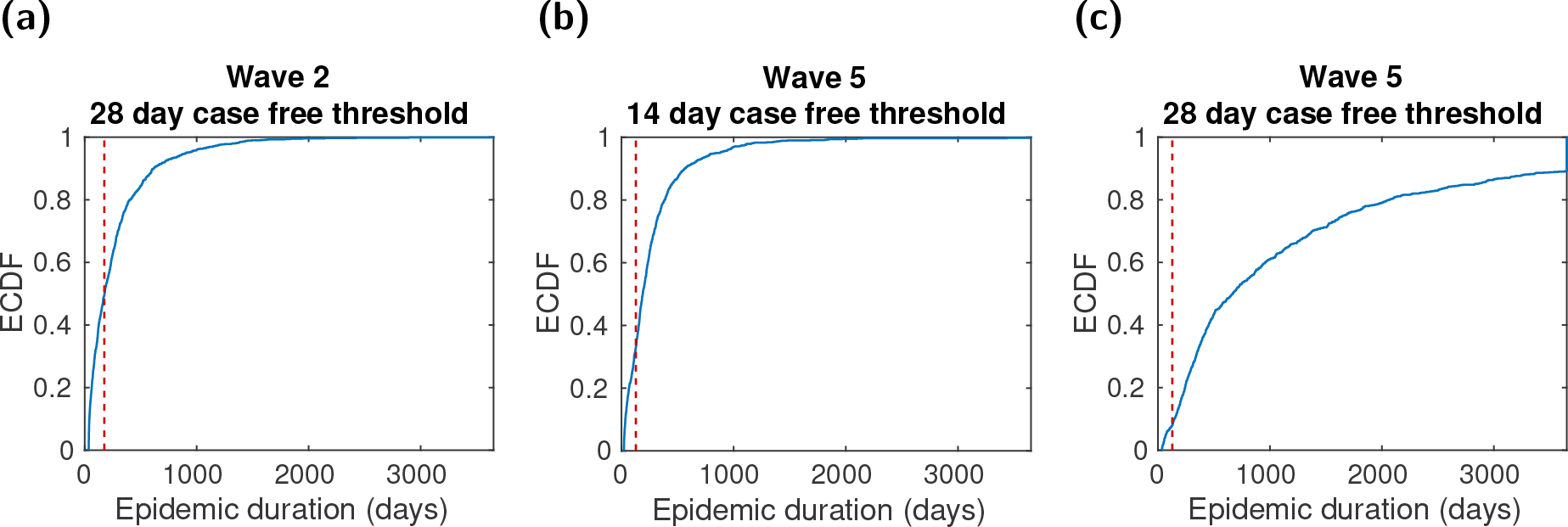
ECDF for epidemic duration from simulations of the specified transmission model, with the given number of consecutive infection-free days required for an outbreak to be deemed as completed. All simulations used infected premises culling only (no additional controls were in place), with reporting to culling times weighted by the empirical probability mass function. The following ECDFs were constructed using 1,000 simulated realisations: **(a)** Wave 2, 28 day threshold value; **(b)** wave 5, 14 day threshold value; **(c)** wave 5, 28 day threshold value. The threshold values for number of infection-free days signifying the end of an outbreak were subsequently set to 28 days and 14 days for runs with the wave 2 and 5 fitted models respectively.

For each poultry-targeted management action, we performed 1,000 simulation runs with the wave 2 fitted transmission model and 500 simulation runs with the wave 5 fitted transmission model. To aid intervention comparisons across the runs, we again used a consistent set of sampled parameter values and initial seed infection locations. The control objectives of interest in this scenario were again focused on outbreak length and size, in particular either increasing the chance of an outbreak being short, maximising the likelihood of an outbreak remaining below a specified size, or minimising the number of poultry destroyed as a result of culling. The particular summary statistics that we therefore chose for these control objectives were as follows: (i) outbreak duration *t* being 90 days or less, (ii) outbreak size *I* not exceeding 25 infected premises, (iii) mean number of poultry culled. We also performed a univariate sensitivity analysis on two vaccination-specific variables, namely vaccine efficacy and effectiveness delay, encompassing ranges of 50-90% for vaccine efficacy and 4-14 days for the effectiveness delay respectively.

## Results

### Choice of control policy based on outbreak origin

Here we consider management of outbreaks whose sole viable route of transmission is premises-to-premises. We establish the severity of control or type of surveillance policy that could be implemented to minimise epidemic duration or probability of a widespread outbreak, dependent upon the district of outbreak origin and capacity constraints.

### Culling and vaccination

In the event of outbreaks with wave 2 type transmission dynamics, regardless of the district of introduction, for minimising the epidemic probability we observe that the optimal ring cull radius increases under less restrictive capacity constraints (Fig. 3(a)). If capacities are low, then 1km-3km radius ring culling was found to be optimal for most districts (Fig. 3(a), left panel). As capacities increase, we observe a slight increase in the optimum radius, with 8-10km ring culling optimal for outbreaks occurring in some districts (Fig. 3(a), right panel).

**Fig. 3.**
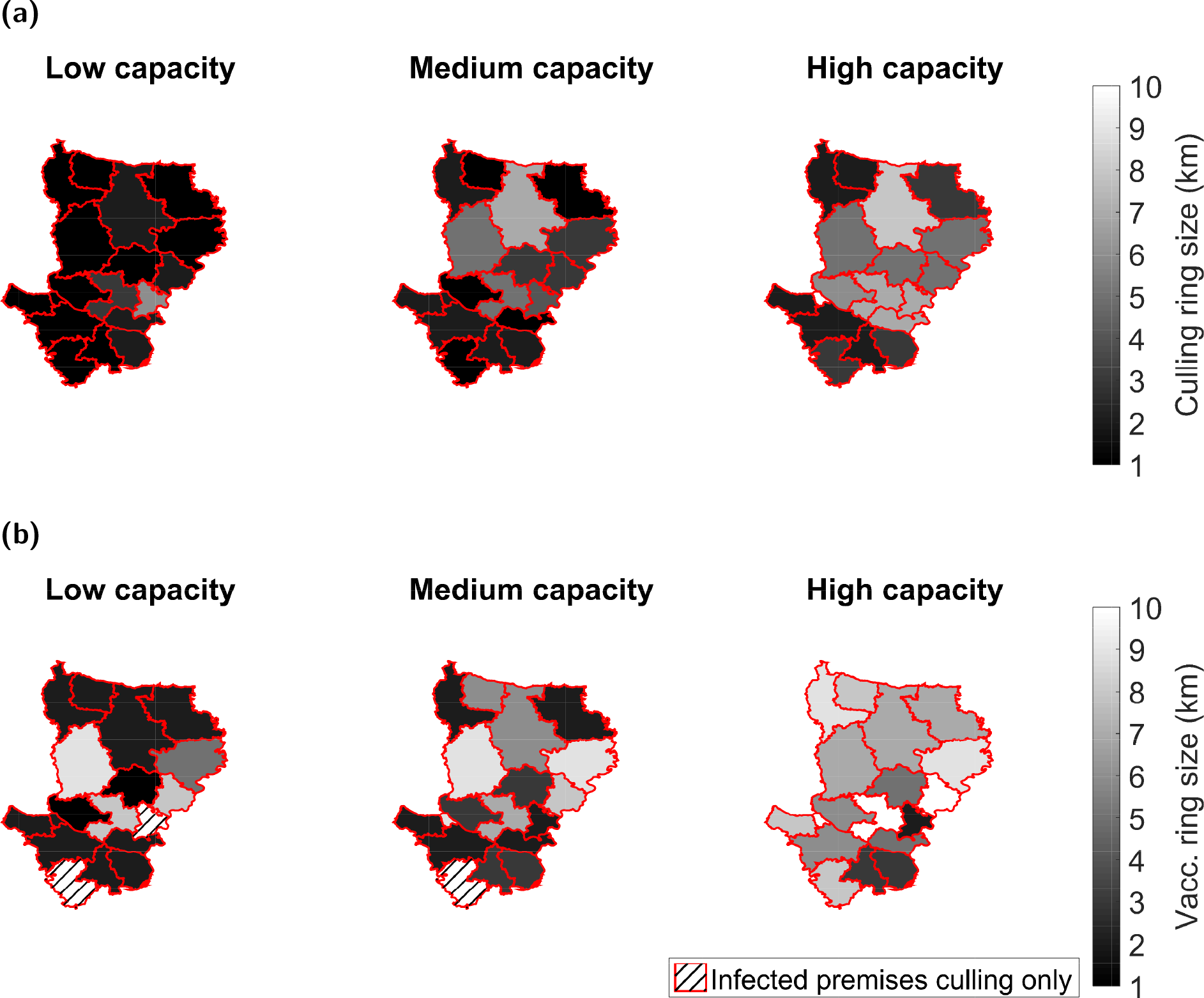
Maps displaying the ring range that optimises minimising epidemic probability with respect to district of outbreak origin and control capacity level, under wave 2 type transmission dynamics. For each combination of control capacity level, district of outbreak origin and control type 1,000 simulation runs were performed. Hatching of a district indicates the preferred strategy was culling infected premises only, while solid shading corresponds to the ring size determined as the optimal severity of response against outbreaks that originally emerged in that district. Lighter shading corresponds to a larger ring culling region. Types of control tested were **(a)** ring culling, and **(b)** ring vaccination. For full results see Table S3.

A similar effect was observed when considering vaccination as a control strategy (Fig. 3(b)). However, for some districts, in conjunction with low and mid-level vaccination capacities, vaccination was not found to decrease the probability of epidemic take off, with solely culling those premises reporting infection the preferred strategy (Fig. 3(b), left and middle panels). Optimal vaccination radii for each capacity level were found to be larger than optimal ring culling radii, possibly owing to a delay in onset of immunity. Qualitatively similar outcomes were observed across the tested transmission models and capacity constraints when the objective was to minimise expected outbreak duration (Figs. S2 and S3).

When analysing the impact of control policies to minimise epidemic risk for outbreaks with wave 5 transmission dynamics, we observe a different effect. In this case, optimal ring culling radii were higher than optimal vaccination radii for many districts, even when capacities to implement control were high (Fig. 4). In low capacity circumstances the epidemic source made scant difference to the chosen ring culling size, typically 1km (Fig. 4(a), left panel). This did not hold under a high resource capacity. Outbreaks emerging in central and northern districts typically required upper radius values of 7km or 8km, while the western district of Rajbari (east) required the 10km upper limit of the range of values explored here. In the event of an outbreak beginning in one of the remaining districts, only localised ring culling of 1km or 2km was suggested, though we observed a ring cull of some form was always found to be preferred over merely culling infected premises (Fig. 4(a), right panel).

**Fig. 4.**
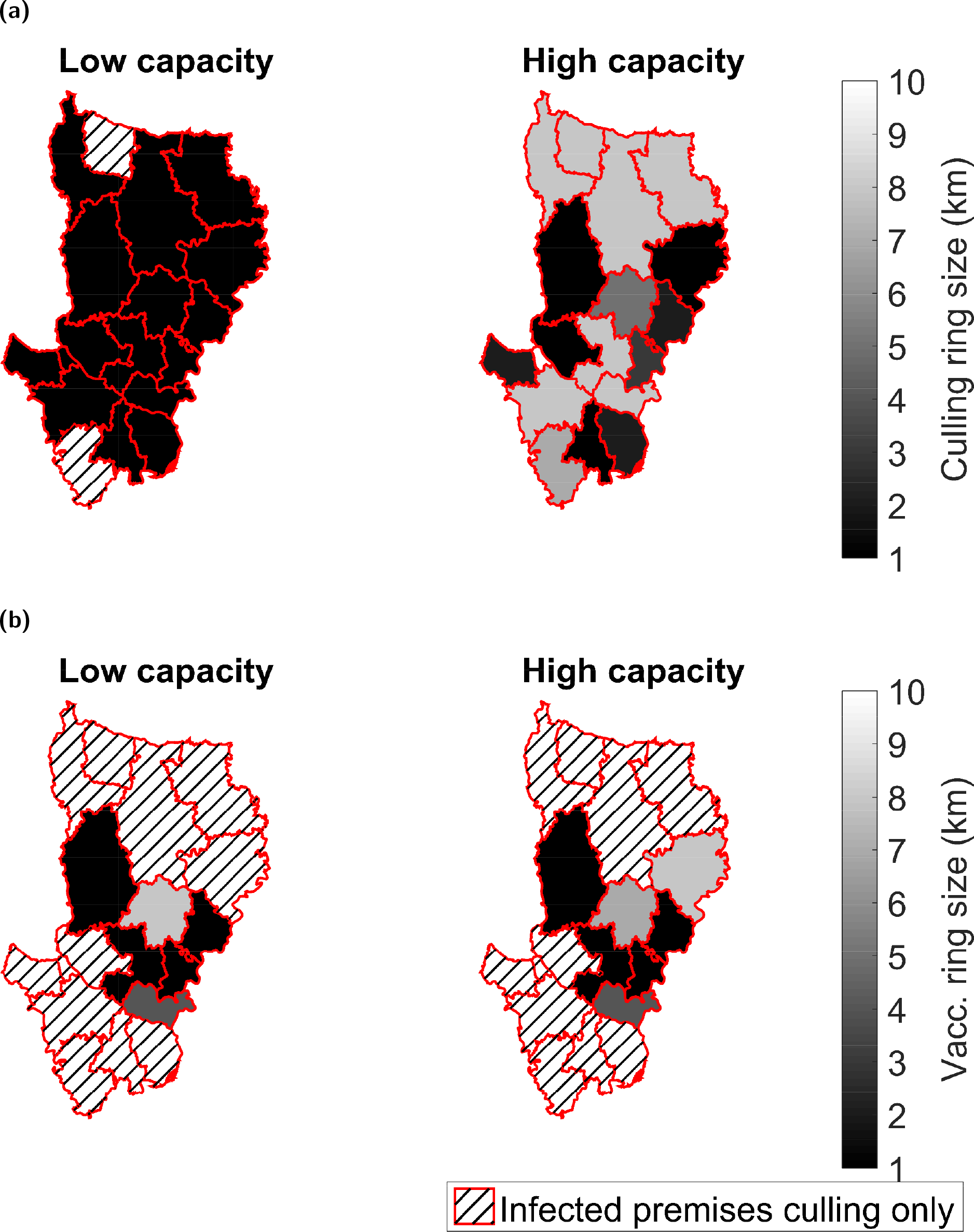
Maps displaying the ring range that optimises minimising epidemic probability with respect to district of outbreak origin and control capacity level, under wave 5 type transmission dynamics. For each combination of intervention method and district of outbreak origin 1,000 simulation runs were performed. Hatching of a district indicates the preferred strategy was culling infected premises only, while solid shading corresponds to the ring size determined as the optimal response against outbreaks that originally emerged in that district. Lighter shading corresponds to a larger intervention region. Types of control tested were **(a)** ring culling, and **(b)** ring vaccination. For full results see Table S5.

On the other hand, regardless of capacity constraints, for outbreaks beginning in northern and southern districts ring vaccination did not provide improved impact over solely culling infected premises, while central districts typically only required a coverage radius of 5km or less (Fig. 4(b)).

As a cautionary note, sensitivity analysis of the variations in the control objective metrics against intervention severity (for outbreaks beginning in a given district) revealed these variations to be small, especially under vaccination measures (Figs. S4 to S7).

### Active surveillance

We now investigate the extent to which H5N1 outbreak burden in the Dhaka division of Bangladesh may be reduced through active surveillance. As described above, we consider implementation of both proactive and reactive surveillance strategies. Our model indicates that, regardless of outbreak wave and location of outbreak, proactive surveillance schemes were optimal across all capacity scenarios and control objectives. Additionally, independent of the source district for the outbreak, higher capacity thresholds usually led to greater reductions in outbreak length and size relative to the scenario where no active surveillance scheme was used (Fig. 5 and Fig. S8).

**Fig. 5.**
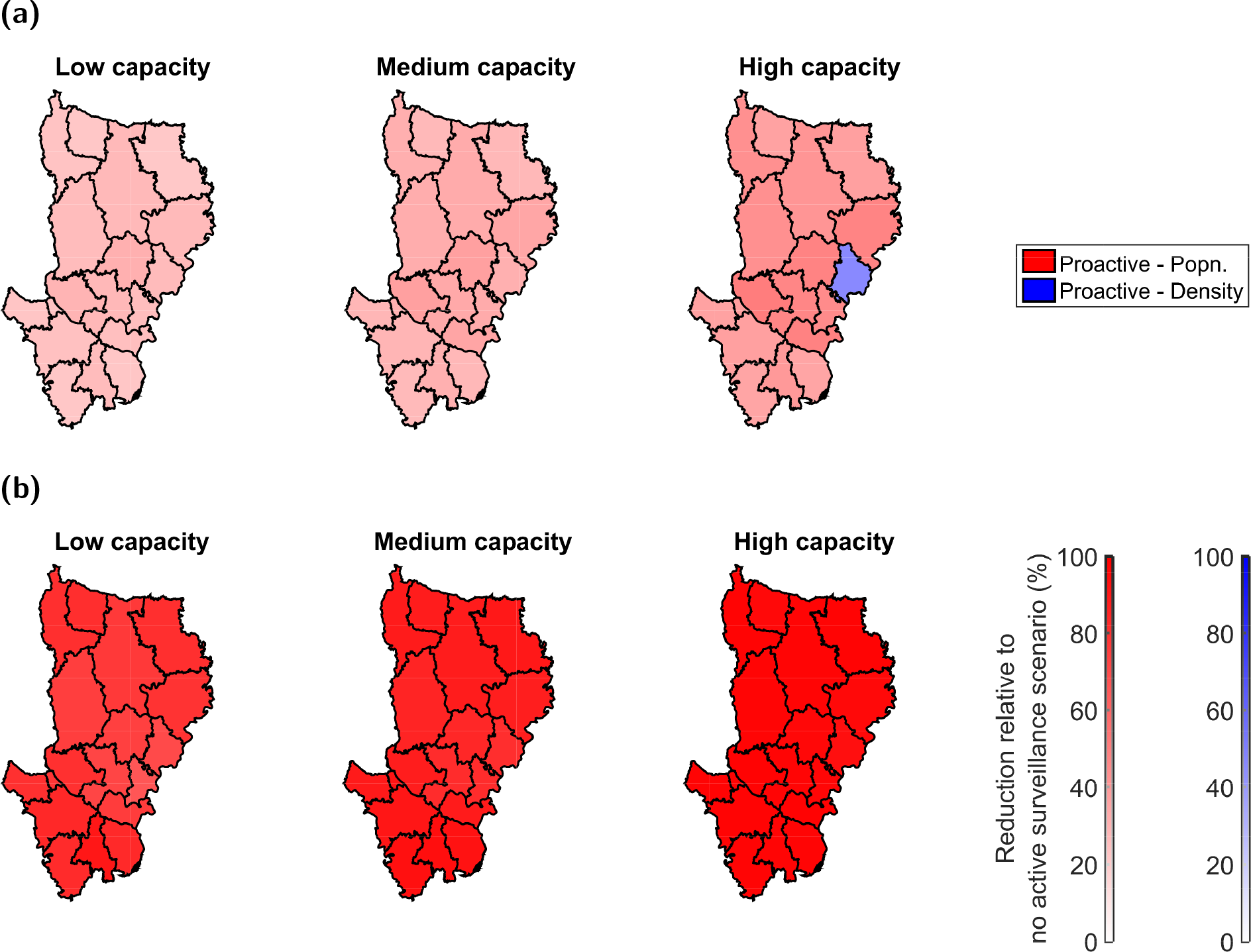
Maps displaying the preferred active surveillance strategy to optimise control objectives with respect to district of outbreak origin and capacity setting, for outbreaks with wave 2 type transmission dynamics. For each combination of active surveillance method and district of outbreak origin 1,000 simulation runs were performed. District colour corresponds to the active surveillance strategy determined to be optimal for countering outbreaks originating from that district (red - ‘proactive by population’, blue - ‘proactive by premises density’). In each case the two reactive schemes, ‘reactive by distance’ and ‘reactive by population’, were also tested, but neither were ever deemed to be the optimal course of action. Transparency coincides with the reduction in the objective metric relative to the scenario where no active surveillance was used, with completely transparent corresponding to a 0% reduction (no improvement) and completely opaque corresponding to a 100% reduction. **(a)** Minimising average outbreak duration control objective - ‘proactive by population’ scheme was generally preferred, although we found discrepancies in the best scheme dependent upon the control capacity setting. **(b)** Minimising the probability of an epidemic control objective - ‘proactive by population’ scheme was found to be preferred in all cases when optimising for this aim. For full results see Tables S7 and S8.

For wave 2 transmission dynamics, the ‘proactive by population’ surveillance strategy was found to be optimal for all capacities and districts, with the exception of the district of Narshingdi where the capacity for active surveillance implementation is high. In this instance, if we are interested in minimising outbreak duration, ‘proactive by premises density’ surveillance could be implemented, whilst ‘proactive by population’ surveillance could be used if we wish to minimise the likelihood of an epidemic (Fig. 5(a) and Fig. 5(b), right panels). Similar outcomes were obtained for outbreaks with wave 5 type transmission dynamics where, irrespective of the district where the outbreak originated, the ‘proactive by population’ strategy was always selected as the optimal action (Fig. S8).

### Cross-intervention performance comparison

For each combination of transmission model, capacity-level and control objective, we compared the top performing strategy within each intervention type (ring culling, ring vaccination, active surveillance) relative to culling infected premises alone. In all circumstances, the best performing active surveillance scheme was deemed to be the preferred approach in optimising the control objectives of interest (Tables S3 to S6).

Particularly noteworthy are the stark reductions (between 65% to 99%) in the probabilities of an epidemic occurring under wave 2 type transmission dynamics when utilising a ‘proactive by population’ active surveillance scheme versus solely culling infected premises. On the other hand, the attained reductions in the expected outbreak duration were generally between 20-50%, thus less prominent (Fig. 5 and Tables S3 and S4). Under wave 5 type transmission dynamics, reductions in the measures for assessing both epidemic duration and epidemic probability control objectives lay in the range of 30-85% (Fig. S8 and Tables S5 and S6).

### Choice of control policy in presence of external factors

In this section, we consider the impact of control in the Dhaka division in the event of external introductions of disease from the surrounding divisions. In this instance, we determine the control or surveillance policy that could usefully be implemented across all districts in the division to minimise the epidemic duration, outbreak size or the number of poultry culled.

### Culling and vaccination

For control objectives targeting outbreak length and magnitude, we ascertained that ring culling typically outperformed ring vaccination, with qualitatively similar outcomes acquired for our two distinct transmission models (Figs. 6 and S9). We found that even when vaccination capacity was high, ring culling resulted in a lower likelihood of long duration outbreaks and fewer premises becoming infected.

**Fig. 6.**
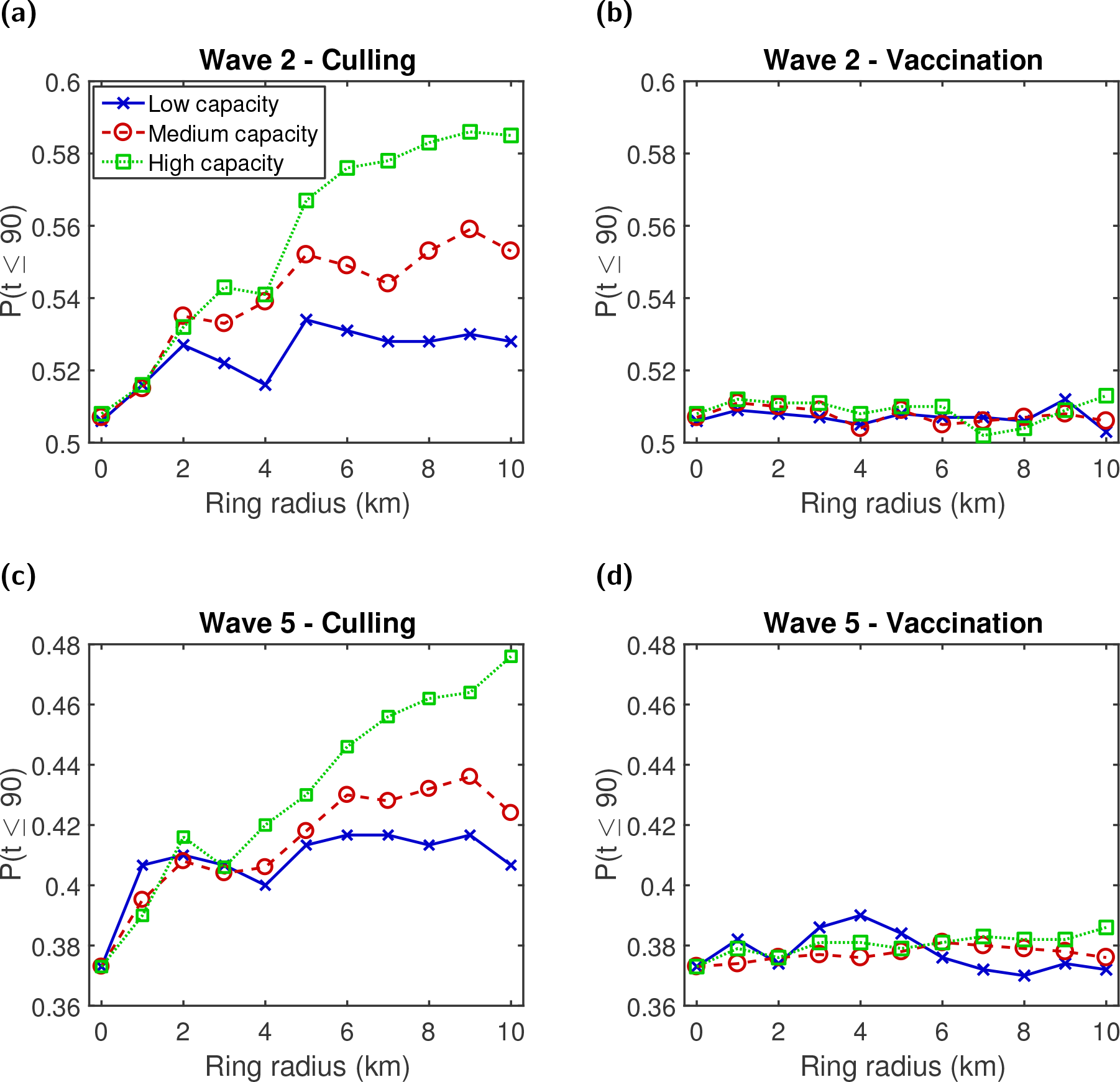
Predicted probability of outbreak duration (t) being 90 days or less for different ring culling and vaccination radii. For each transmission model and control method combination, the three capacity settings of interest, low (solid blue line, crosses), medium (dashed red line, circles), and high (dotted green line, squares) displayed disparate behaviour. **(a)** Wave 2 - culling; **(b)** wave 2 - vaccination; **(c)** wave 5 - culling; **(d)** wave 5 - vaccination. In all panels a ring size of 0km corresponds to a control action of culling infected premises only. Results are averaged over 1,000 simulations and 500 simulations for wave 2 and wave 5 type transmission dynamics respectively.

For ring culling there was evidence of a performance hierarchy across the three tested capacity constraints Figs. 6(a) and 6(c). For any given ring size, a high capacity allowance generally outperformed a medium capacity allowance, which in turn outperformed a low capacity allowance. Further, under high control capacity resource availability, each incremental increase in the radius size generally led to modest improvements in the summary output of interest (at least up to the 10km upper limit in place here). In contrast, for low and medium capacity thresholds, the optimal radius size varied dependent upon the objective of interest. Such a relationship was less apparent for vaccination. For the epidemic duration control metric, irrespective of the transmission dynamics, we identified little variation in this measure among the three capacity constraints across all tested ring sizes and also relative to only culling infected premises Figs. 6(b) and 6(d). Comparable outcomes were found when optimising the epidemic size objective of *I* ≤ 25 (Fig. S9). The epidemic duration and size measures were both insensitive to a range of vaccine efficacies and vaccine effectiveness delay times (Figs. S10 and S11).

However, if our objective was to minimise the total number of poultry culled, we found that vaccination was, unsurprisingly, preferred over ring culling in all instances (Fig. 7). Incremental increases in vaccination radius size under each set of control capacity conditions were found to cause modest improvements with regard to this objective. Specifically, a 9km or 10km ring was optimal across all capacities and both transmission models. On the other hand, if conditions preclude the use of vaccination, pursuing a ring culling strategy in combination with this control objective results in the best performing action being either no culling beyond infected premises or a ring cull of 1km (Fig. 7). Under wave 2 type transmission dynamics, high capacity ring culling results in the largest number of poultry culled, particularly when implemented at large radii (Fig. 7(a)). For wave 5 transmission dynamics the opposite effect is seen (Fig. 7(c)). The larger expected size of outbreaks under wave 5 type transmission dynamics (relative to wave 2 type transmission dynamics) means that low capacity ring culling proves insufficient to control the outbreak, resulting in a much larger number of poultry being culled than for higher capacities. For either wave 2 and wave 5 type transmission, increasing vaccine efficacy or decreasing the vaccine effectiveness delay led to modest reductions in the expected number of poultry culled per outbreak (Figs. S10 and S11).

**Fig. 7.**
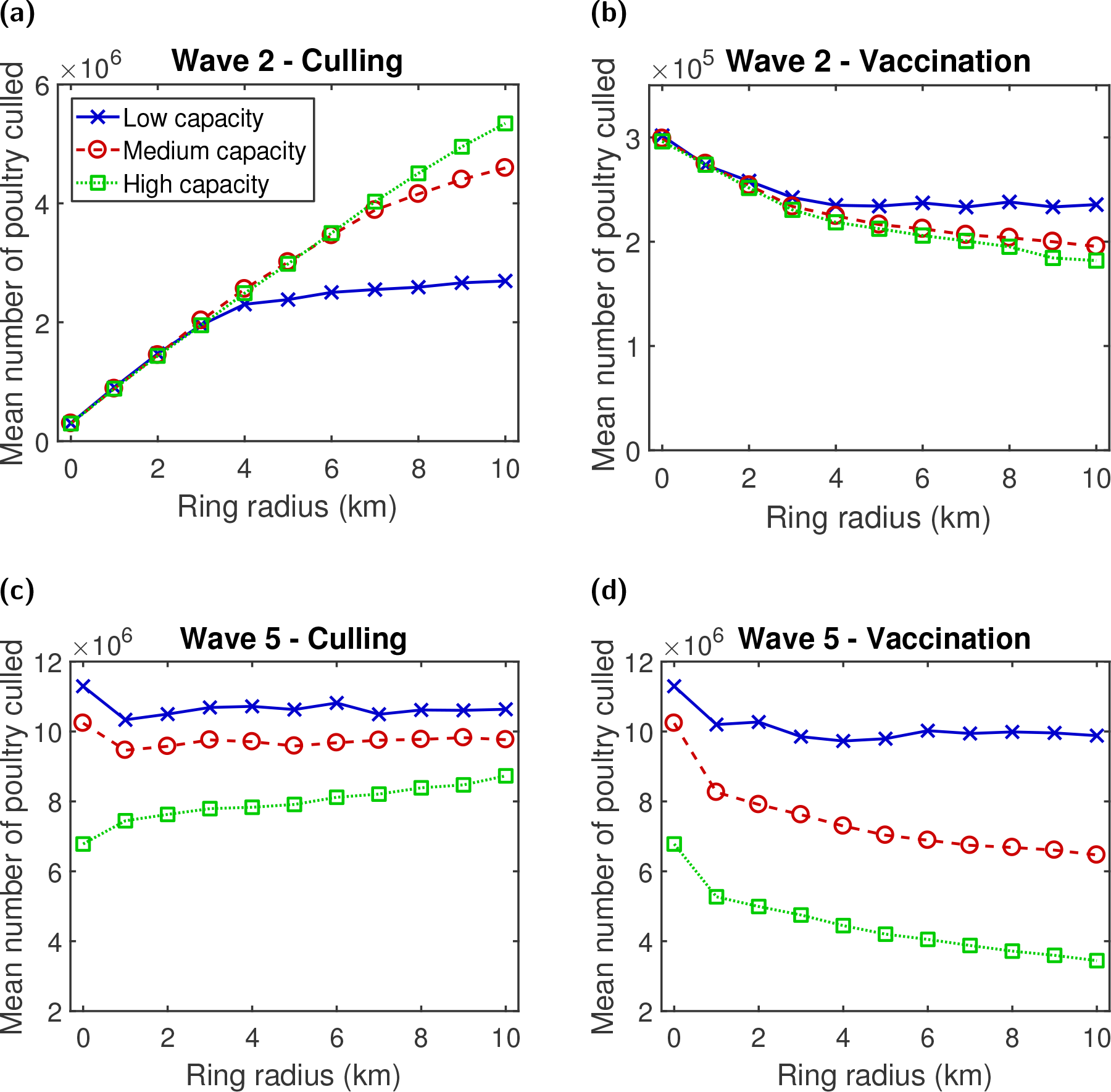
Mean number of poultry culled for different ring culling and vaccination radii. The three capacity settings of interest were low (solid blue line, crosses), medium (dashed red line, circles), and high (dotted green line, squares). If pursuing a ring culling strategy, either no culling beyond infected premises or a ring cull of 1km were deemed optimal. For a ring vaccination strategy, a 9km or 10km ring was selected across all capacities. **(a)** Wave 2 - culling; **(b)** wave 2 - vaccination; **(c)** wave 5 - culling; **(d)** wave 5 - vaccination. In all panels a ring size of 0km corresponds to a control action of culling infected premises only. Results are averaged over 1,000 simulations and 500 simulations for wave 2 and wave 5 type transmission dynamics respectively.

### Active surveillance

Investigating the effectiveness of active surveillance against H5N1 HPAI under this transmission setting, a collection of common trends were obtained across the three control objectives (outbreak duration being 90 days or less, outbreak size not exceeding 25 premises, minimising mean number of poultry culled) and two disease transmission models analysed.

Irrespective of the objective being scrutinised, the most effective active surveillance policy was the ‘proactive by population’ scheme, with this conclusion being consistent under either wave 2 or wave 5 type transmission dynamics (Figs. 8(a) to 8(c)). Additionally, increased availability of resources for control raised the performance of this kind of action. This is typified when examining the outbreak duration objective of *t* ≤ 90. Under the wave 2 transmission model this rose from 0.55 (low capacity) to 0.61 (high capacity), whereas with no active surveillance in use the probability was only 0.51. Such effects were even more stark for the wave 5 transmission model, with outbreaks being more likely to spread rapidly and having enhanced longevity. With an initial value of 0.38 when no active surveillance was used, this rose to 0.46 for low capacity levels, reaching 0.58 under high capacity conditions. Thus, use of the wave 5 transmission model led to an approximate 50% improvement over having no control.

**Fig. 8.**
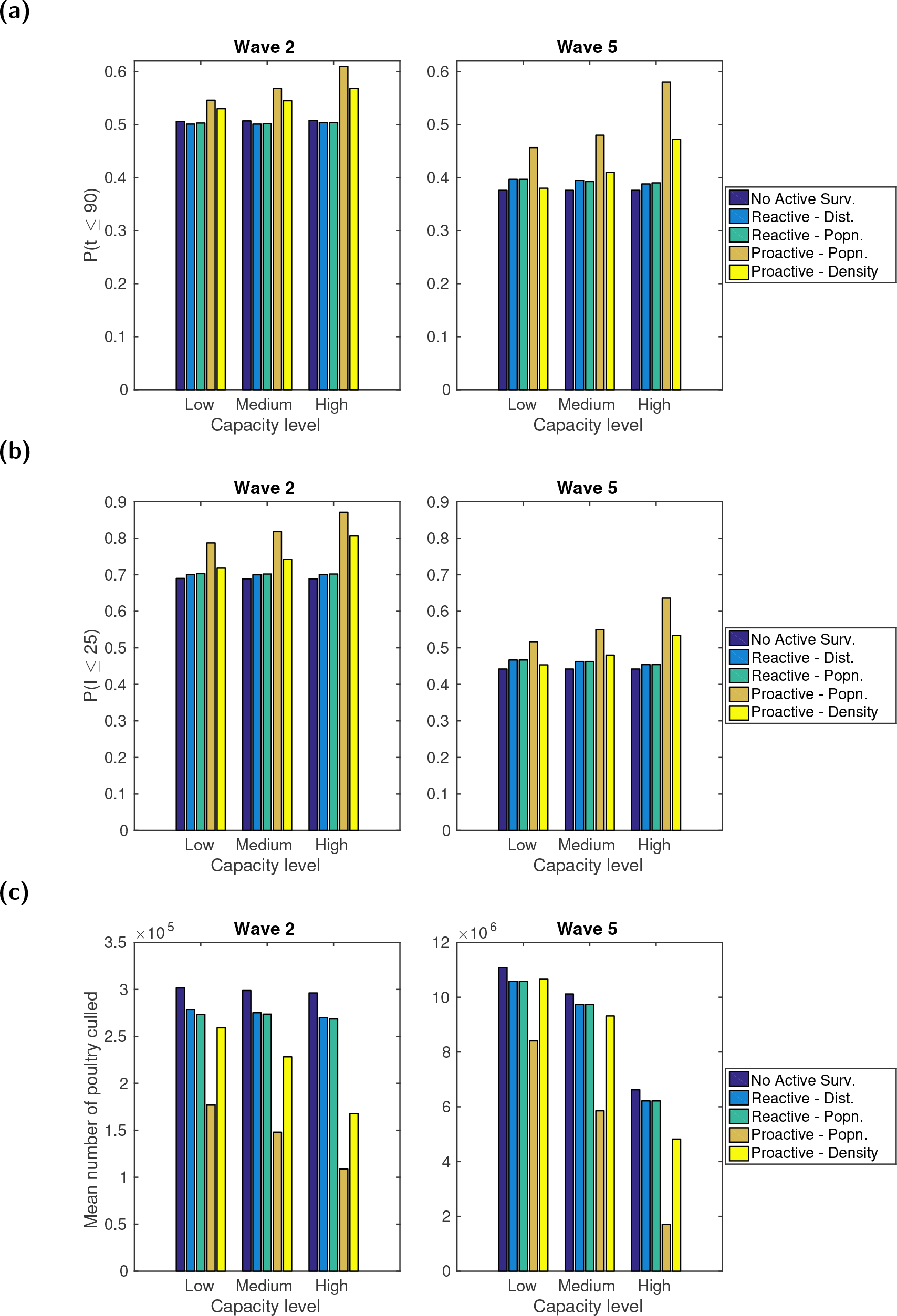
Bar plots comparing the impact of different active surveillance strategies on specific control objectives. For each combination of transmission model, resource restrictions and active surveillance strategy we performed between 500 and 1,000 simulation runs. The control objectives were: **(a)** predicted probability for outbreak duration *t* being 90 days or less; **(b)** predicted probability for outbreak size *I* not exceeding 25 premises; **(c)** mean number of poultry culled. For both wave 2 and wave 5 transmission dynamics the ‘proactive by population’ surveillance strategy was found to be optimal for all control objectives considered, irrespective of the capacity limitations. Full values are given in Tables S11 to S13

Although the ‘proactive by premises density’ strategy offers notable improvements under less stringent capacity constraints, it was not as effective as the population-based targeting measure. This is exemplified by the discrepancy between the two typically growing with enlarged capacity thresholds. For example, the difference grew from 0.02 (at low capacity) to 0.04 (at high capacity) for *t* ≤ 90 using the wave 2 transmission model, and from 0.07 (at low capacity) to 0.11 (at high capacity) for *I* ≤ 25 using the wave 5 transmission model. A further drawback of the ‘proactive by premises density’ strategy was that under low control capacity levels it struggled to beat either reactive surveillance policy (Fig. 8).

Comparing the two reactive strategies we found their performance differential to be minor. Despite offering marginal benefits over having no active surveillance policy in use, they did not bring about noticeable improvements towards the desired goal under more relaxed capacity constraints (Fig. 8). The observation of ‘proactive by population’ outperforming ‘proactive by premises density’, and the two reactive strategies only being a slight improvement compared to having no active surveillance, is also evident when comparing the complete premises outbreak size distributions (Fig. S12). For a full listing of values related to the features raised see Tables S11 to S13.

### Cross-intervention performance comparison

In a similar manner to when we previously compared intervention types when optimising control policy based on outbreak origin, we again examine the top performing strategy within each intervention type (ring culling, ring vaccination, active surveillance) relative to culling infected premises alone. Once more, this illuminated the superior performance of ring culling to ring vaccination when aiming to optimise the outbreak size and duration control objectives considered here, and vice versa if optimising the poultry culled control objective (Fig. 9).

**Fig. 9.**
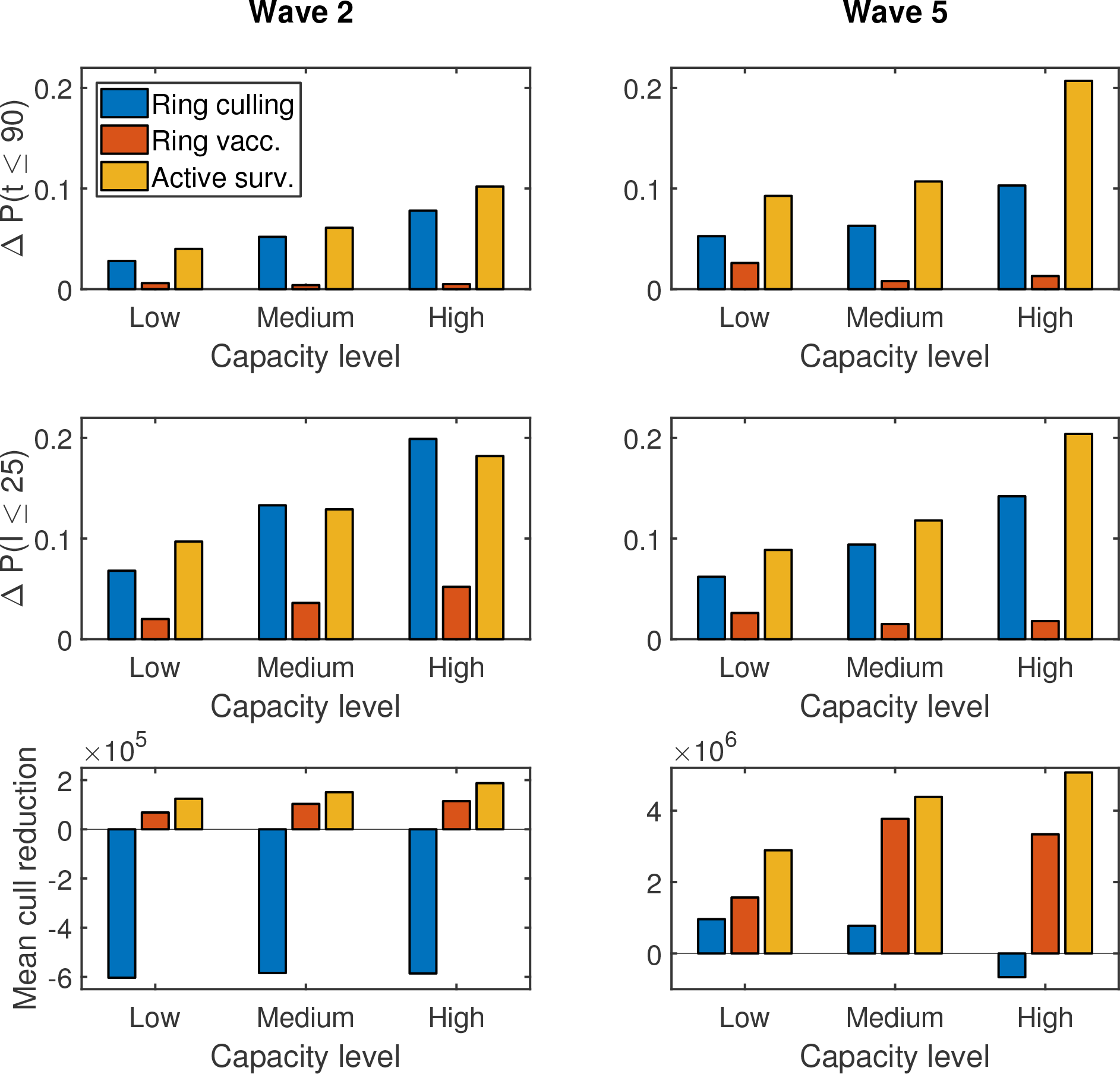
Cross-intervention performance comparison, relative to only culling infected premises. For each combination of transmission model, capacity-level and control objective, we compared the top performing strategy within each intervention type relative to culling infected premises alone. In each panel, the bar order is as follows: Ring culling (bar one, blue), ring vaccination (bar two, orange), active surveillance (bar three). The control objective comparisons were: **(row one)** improvement in predicted probability for outbreak duration *t* being 90 days or less; **(row two)** improvement in predicted probability for outbreak size *I* not exceeding 25 premises; **(row three)** reduction in mean number of poultry culled. Transmission dynamics: **(column one)** wave 2; **(column two)** wave 5. For the majority of scenarios, active surveillance was the dominant strategy.

Active surveillance, in the form of the ‘proactive by population’ scheme, was dominant in the majority of scenarios over the entire range of ring cull and vaccination severities. The exception to this was under wave 2 transmission dynamics when wanting to minimise the probability of the outbreak size exceeding 25 premises, in tandem with there being less constraints on control capacity. Specifically, under medium and high capacity conditions a 9km ring cull was predicted to give the greatest gains relative to only culling infected premises (Fig. 9).

## Discussion

This study explores the predicted repercussions of a variety of intervention methods aimed at the commercial poultry sector within the Dhaka division of Bangladesh, namely culling, vaccination and active surveillance, for mitigating the impact of H5N1 HPAI outbreaks; evaluations were carried out under a scenario where the commercial poultry premises in the region initially began free of H5N1 HPAI viruses. Informed via a mathematical and computational approach, it emphasises how knowledge of both disease transmission dynamics and potential resource limitations for implementing an intervention can alter what are deemed the most effective actions for optimising specific H5N1 influenza control objectives. Likewise, we saw differences in policy recommendations when comparing alternative control objectives to one another. This corroborates previous work that showed establishing the objective to be optimised is pivotal in discerning the management action that should be enacted [17], whilst underlining the potential pivotal role mathematical modelling has in providing decision support on such matters.

A consistent outcome across all combinations of transmission model, capacity constraints and control objectives was the superior performance of proactive schemes, which constantly monitor a predetermined set of premises based on selective criteria, over reactive surveillance schemes (only enforced once an outbreak has begun), ring culling and ring vaccination. Out of the tested proactive schemes, we discerned that monitoring premises with the largest flocks was the most effective approach, with larger coverage levels strengthening performance outcomes. The abovementioned conclusions are further strengthened by being maintained irrespective of the spatial origin of the outbreak.

This may lead us to posit proactive surveillance measures, with flock size based prioritisation, being superior to other control initiatives when applied in other settings. As a note of caution, generalising such guidance to alternative circumstances first necessitates gleaning similar outcomes when applying the methodology to other datasets and spatial scenarios. A caveat of our modelling framework is that potential reporting biases between premises have yet to be considered, as a consequence of discrepancies in the enforcement of biosecurity protocols for example. If premises with larger flocks were to have tighter biosecurity protocols, potentially reducing the standard notification delay relative to other premises (i.e. below seven days), this may curtail the performance of population-based surveillance measures. On the other hand, this highlights an alternative application of this particular methodology, where we instead interpret the reduction in the notification delay as capturing an inherent reporting bias linked to a specific risk factor (such as flock size). Nevertheless, in the instance of Dhaka division in Bangladesh, we have revealed the potential value attached to establishing systems that ensure premises flock size data are both reliable and frequently updated.

Prioritisation schemes linked to flock size would be the most challenging to implement out of those considered in this study, with premises poultry populations fluctuating over time. This is exemplified by the ad hoc approach to collecting commercial poultry premises information via census in Bangladesh, last done in 2010. Nonetheless, such information is theoretically attainable. Expressly for Bangladesh, efforts to monitor commercial poultry flock sizes are facilitated by the ‘Animal Diseases Act 2005’, requiring all commercial poultry premises to be registered. Presently, listings of premises will be maintained in local administration level systems. Although not yet contained in a centralised database, efforts are being made by the Bangladesh DLS to fulfil such a plan (M.G. Osmani and M.A. Kalam, personal communication). This would provide a platform with the capability of receiving revised poultry flock sizes with greater regularity.

Though putting an active surveillance system into action may face difficulties, typified by the previous active surveillance system in Bangladesh being discontinued in 2013 for monetary reasons (M.G. Osmani, personal communication), there are other ongoing surveillance schemes within the country demonstrating the capability to carry out policies of this nature. One example is environmental sampling being used to monitor the situation within LBMs [32]. Another is FAO supported trial surveillance programs, comprising villages being surveyed twice a week and deploying rapid detection tests if HPAI viruses are suspected. With the necessary resources, the existence of these ongoing schemes offers a foundation for the introduction of a larger-scale premises-focused surveillance system (M.A. Kalam, personal communication).

Surveillance methodology is a discipline requiring greater attention. In the context of early detection of the introduction and spread of H5N1 HPAI viruses, active surveillance does not have to be restricted to only looking for clinical signs of disease within poultry flocks. Sustained swabbing and testing of blood samples on targeted premises may allow near real-time detection of viral infections, thereby further minimising the reporting delay, or even fully eradicating it. Other usages of active surveillance include tracing the likely chain of transmission, overseeing poultry value chains involving different poultry products (i.e. the full range of activities required to bring poultry products to final consumers) to ascertain if there is a particular section of the system where biosecurity is compromised, and monitoring trade and marketing links to track the genetic diversity of circulating strains [33, 34]. Such endeavours will in turn contribute towards the standardisation of sampling, testing, and reporting methods, bolstering full-genome sequencing efforts and encouraging sharing of isolates with the scientific community [35].

Notwithstanding the outcomes relative to active surveillance, our assessment of ring culling and ring vaccination unveiled insights into the suggested ring radii sizes if pursuing these classical intervention methods.

For circumstances where transmission is exclusively premises-to-premises, we found considerable variation in the preferred control strategy depending upon the spatial location of the source of the outbreak, the relationship between risk of transmission and between premises distance (examined here by comparing the wave 2 and wave 5 transmission models), and the capacity restrictions that are in place. Although there was a common trend of increasing the suggested radius of an intervention ring zone for less stringent capacity settings, solely culling infected premises was sometimes expected to be the best course of action when both vaccination and ring culling were considered. This is strongly exhibited in the case of reducing the likelihood of a widespread outbreak for an infection with wave 5 type transmission dynamics, with additional ring vaccination deemed ineffective for the majority of district origin locations. In such cases, it may therefore be necessary to consider alternative intervention measures other than vaccination, such as strict movement controls, to further reduce the risk of disease spread. The robustness of these outcomes to alternate vaccine efficacies and the assumptions of effectiveness delay merits further investigation. Given insight into the exact outbreak circumstances, the abundant variation in the preferred control action dependent upon the origin of the outbreak shows the potential benefits of having flexibility to adapt the intervention that is ratified.

Under situations where external factors have a meaningful impact on the transmission dynamics, we found that the class of intervention preferred was highly dependent upon the objective of the control policy. If we are interested in either minimising outbreak duration or the number of infected premises, ring culling is preferred to vaccination. Finding that ring culling outperforms ring vaccination may be a result of the vaccine assumptions, a seven day delay from vaccination to immunity and a 70% vaccine efficacy, though qualitative conclusions were unaltered when analysing sensitivity to these vaccine-specific variables. If minimising the number of poultry culled is a priority, then ring vaccination is naturally preferred over ring culling. Furthermore, we observe effects of capacity becoming apparent for vaccination rings of over 4km, as limited capacity interventions applied beyond this rather local scale did not demonstrate additional increases in effectiveness. Situations may arise where ring culling is used in conjunction with this control objective, chiefly when vaccination is not an intervention choice. In such circumstances, one might expect no culling beyond infected premises to be deemed the best action, regardless of the invoked capacity constraints and the underlying transmission dynamics. Nevertheless, highly localised ring culls of 1km were preferred in some instances.

It is vital that the area covered by ring based control methods is selected to only be as large as necessary. If set too small then other premises just outside the intervention zone may become infected, which would have been contained had harsher measures been imposed. However, the use of widespread pre-emptive culling based on defined areas around an outbreak has been shown to be very difficult to implement effectively in developing countries. Enforcing wide area culling can alienate farmers if healthy birds are destroyed and the reimbursement through compensation is deemed inadequate or is provided too late. Loss of poultry owner cooperation can be counterproductive, leading to resentment and resistance to further control measures [21].

An alternative focal point for control, not explicitly included here, is trade and LBMs. In the event of disease outbreaks among poultry, both farmers and traders face economic losses. In order to reduce such loss they may modify their practices, altering the structure of the trade networks. Reshaping the trade network may in turn modify the disease transmission dynamics and possibly facilitate additional spread [36].

The high density and variety of avian hosts in Bangladeshi LBMs supports the maintenance, amplification and dissemination of avian influenza viruses, whilst providing frequent opportunities for inter-species transmission events [37, 38]. In a meta-analysis of before-after studies, to assess the impact of LBM interventions on circulation of avian influenza viruses in LBMs and transmission potential to humans, Offeddu et al. [39] determined that periodic rest days, overnight depopulation and sale bans of certain bird species significantly reduced the circulation of avian influenza viruses in LBMs. Furthermore, prolonged LBM closure reduced bird-to-human transmission risk. Developing a theoretical model incorporating LBMs and trade networks would allow us to validate these previous findings.

The analysis presented here did not consider the role of domestic ducks, due to the low number of poultry premises within the Dhaka division recorded as having ducks present. Nonetheless, at a national level domestic ducks are part of an intricate animal production and movement system, which may contribute to avian influenza persistence [40]. Ducks raised in free-range duck farms in wetland areas have considerable contact with wild migratory birds in production sites, and subsequently with other poultry animals in LBMs. Furthermore, influenza viruses of the H5 subtype typically persist in ducks with very mild or no clinical signs [41–45], affecting epidemic duration and spread. If applying this work to other regions of Bangladesh, or scaling it up to encompass the entire country, domestic ducks warrant inclusion within the model framework.

This initial analysis can be extended naturally in a number of additional ways to those already mentioned. While we considered conventional control strategies used to combat avian influenza outbreaks among poultry, namely culling, vaccination and active surveillance, one could compare these traditional schemes with innovative direct interruption strategies that modify the poultry production system [46]. An example would be intermittent government purchase plans, so that farms can be poultry-free for a short time and undergo disinfection. Another is to model restrictions on species composition. This aims to synchronise all flocks on a premises to the same birth-to-market schedule, allowing for disinfection of the premises between flocks. A separate direction for further study is to understand whether the intensification of farming systems, which can alter the demography and spatial configuration of flocks, requires the severity of previously established control protocols to be amended to prevent a small-scale outbreak developing into a widespread epidemic. Such an analysis may be realised by modifying the current model framework to classify premises based on flock size and whether they use intensive or extensive methods, with distinct epidemiological parameters for each group.

The extent to which other premises prioritisation schemes for administering the intervention of interest influences the results also warrants further examination. For the culling and vaccination controls deliberated here we assumed premises were prioritised by distance, from the outer edge of the designated ring control size inwards. Alternative prioritisation strategies that may be considered, subject to availability of the necessary data, include ordering by flock size (in either ascending or descending order), by between-premises flock movement intensity or prioritising by value chain networks. In the case of active surveillance, rather than a fixed, pre-determined policy, extra flexibility can be included by allowing for differing pre- and post-outbreak strategies. Ultimately, public-health decision making generally necessitates the real-time synthesis and evaluation of incoming data. Optimal decision making for management of epidemiological systems is often hampered by considerable uncertainty, with epidemic management practices generally not incorporating real-time information into ongoing decision making in any formal, objective way. An adaptive management approach could be implemented to account for the value of resolving uncertainty via real-time evaluation of alternative models. In addition, this procedure naturally includes economic models embedded within a mathematical framework, allowing for the assessment of control measures to be undertaken in monetary terms [16, 17].

To conclude, through the use of mathematical modelling and simulation, the results of this paper illustrate some general principles of how disease control strategies directed against H5N1 avian influenza outbreaks amongst (initially H5N1 free) commercial poultry premises in the Dhaka division of Bangladesh could be prioritised and implemented, accounting for both resource availability and the particular control objective being optimised. We highlight how targeting of interventions varies if it is believed transmission is predominately premises-to-premises, versus the scenario where importations and other external factors are included. Based on this consideration, reactive culling and vaccination control policies could beneficially pay close attention to transmission factors to ensure intervention targeting is optimised. Yet, irrespective of disease transmission assumptions, amongst all considered interventions we found proactive surveillance schemes that target sites with the largest poultry flocks to typically be the most impactful in reducing the scale of a developing outbreak of H5N1 avian influenza. Consequently, we advocate that much more attention be directed at identifying ways in which control efforts can be targeted for maximum effect.

## Acknowledgements

We thank the Bangladesh Department of Livestock services (DLS) for providing the premises and live bird market data. Colleagues at FAO-ECTAD (Emergency Centre for Transboundary Animal Diseases) office in Bangladesh are thanked for their contribution. We acknowledge USGS (United States Geological Survey) internal reviewers for providing constructive feedback on the manuscript, plus Matt Keeling and Nick Savill for helpful discussions. The work described in the paper was partially supported by the USAID Emerging Pandemic Threats Program (EPT) and the authors would like to thank them for their continued support. The use of trade, product, or firm names in this publication is for descriptive purposes only and does not imply endorsement by the U.S. Government.

## Author contributions

**Conceptualisation:** Edward M. Hill, Thomas House, Michael J. Tildesley.

**Formal analysis:** Edward M. Hill.

**Investigation:** Edward M. Hill, Wantanee Kalpravidh, Subhash Morzaria, Muzaffar G. Osmani, Eric Brum, Mat Yamage, Md. A. Kalam, Xiangming Xiao.

**Methodology:** Edward M. Hill, Thomas House, Marius Gilbert, Michael J. Tildesley. Software: Edward M. Hill.

**Supervision:** Thomas House, Michael J. Tildesley.

**Visualization:** Edward M. Hill, Michael J. Tildesley.

**Writing - original draft:** Edward M. Hill, Michael J. Tildesley.

**Writing - review & editing:** Edward M. Hill, Thomas House, Madhur S. Dhingra, Md. A. Kalam, Diann J. Prosser, John Y. Takekawa, Xiangming Xiao, Marius Gilbert, Michael J. Tildesley.

## Data availability

Data are available from FAO Regional Office for Asia and the Pacific who may be contacted at

